# Impact of sleep fragmentation, heart failure, and their combination, on the gut microbiome

**DOI:** 10.1101/2020.09.11.294447

**Authors:** Olfat Khannous-Lleiffe, Jesse R. Willis, Ester Saus, Ignacio Cabrera-Aguilera, Isaac Almendros, Ramon Farré, David Gozal, Nuria Farré, Toni Gabaldón

**Affiliations:** Barcelona Supercomputing Centre (BSC-CNS). Jordi Girona, 29. 08034. Barcelona, Spain; Institute for Research in Biomedicine (IRB Barcelona), The Barcelona Institute of Science and Technology, Baldiri Reixac, 10, 08028 Barcelona, Spain; Unitat de Biofísica i Bioenginyeria, Facultat de Medicina i Ciències de la Salut, Universitat de Barcelona, Barcelona, Spain; Department of Human Movement Sciences, Faculty of Health Sciences, School of Kinesiology, Universidad de Talca, Talca, Chile; CIBER de Enfermedades Respiratorias, Madrid, Spain; Institut d’Investigacions Biomèdiques August Pi i Sunyer, Barcelona, Spain; Department of Child Health and Child Health Research Institute, The University of Missouri School of Medicine, Columbia, MO, United States; Heart Failure Unit, Department of Cardiology. Hospital del Mar (Parc de Salut Mar). Barcelona; Heart Diseases Biomedical Research Group, IMIM (Hospital del Mar Medical Research Institute), Barcelona, Spain; Department of Medicine, Universitat Autònoma de Barcelona, Barcelona, Spain; Catalan Institution for Research and Advanced Studies (ICREA), Barcelona, Spain

**Keywords:** Metagenomics, Microbiome, Sleep fragmentation, Heart failure, Sleep apnea

## Abstract

Heart failure (HF) is a common condition associated with a high rate of hospitalizations and adverse outcomes. HF is characterized by impairments of the cardiac ventricular filling and/or ejection of blood capacity. Sleep fragmentation (SF) involves a series of short sleep interruptions that lead to fatigue and contribute to cognitive impairments and dementia. Both conditions are known to be associated with increased inflammation and dysbiosis of the gut microbiota. In the present study, male mice were distributed into four groups, and subjected for four weeks to either HF, SF, both HF and SF, or left unperturbed as controls. We used 16S metabarcoding to assess fecal microbiome composition before and after the experiments. Evidence for distinct alterations in several bacterial groups and an overall decrease in alpha diversity emerged in HF and SF treatment groups. Combined HF and SF conditions, however, showed no synergism, and observed changes were not always additive, suggesting that some of the individual effects of either HF or SF cancel each other out when applied concomitantly.

**IMPORTANCE:** The study demonstrates the potential of the gut microbiome as a source of molecular markers for the diagnosis, prevention, and treatment of both heart failure and sleep fragmentation conditions in isolation. Our results provide the first evidence of an antagonistic effect of the presence of both conditions in the gut microbiome dysbiosis, showing an attenuation of the alterations that are observed when considering them separately.

## INTRODUCTION

Heart failure (HF) is a prevalent disease associated with a poor, yet variable prognosis whose causal mechanisms are not entirely understood (Camps-Vilaro et al. 2020). Comorbidities, such as sleep apnea, are frequent in patients with HF, and have been associated with a worsened prognosis (Farre et al. 2017). The adverse outcomes associated with the co-existence of HF and sleep apnea have been attributed, at least in part, to excessive activation of the sympathetic autonomic nervous system (Cowie et al. 2017, Javaheri et al. 2020), yet there is substantial variability underlying these relationships suggesting that other upstream factors may be also involved. Among these factors, the gut microbiome, a vast and complex polymicrobial community that coexists with the human host and is extraordinarily adaptable to a variety of intrinsic or extrinsic changes, plays an important role in the development of immunological phenotypes and in host metabolism (Tremaroli et al. 2012), and could be implicated in the adverse outcomes of HF-sleep apnea (Mashaqi et al. 2019).

Indeed, previous studies have shown evidence implicating the gut microbiome in the physiopathology and prognosis of HF (Tang et al. 2017). HF is associated with reduced microbiome diversity (Luedde et al. 2017) and a shift in the major bacterial phyla, resulting in a lower Firmicutes/Bacteroidetes ratio (Mayerhofer et al. 2020), an increase in Enterbacterales, *Fusobacterium* and *Ruminococcus gnavus*, but also in a decrease in *Coriobacteriaceae*, *Erysipelotrichaceae, Ruminococcaceae*, and *Lachnospiraceae* (Luedde et al 2017). Moreover, some intestinal microbial metabolites (e.g. trimethylamine-N-oxide (TMAO) and its precursors) are present in higher amounts in patients with chronic HF, and elevated levels of TMAO have been independently associated with an increased risk of mortality in acute and chronic HF (Suzuki et al. 2016). Furthermore, patients with HF, present high blood levels of endotoxins, lipopolysaccharides (LPS), and tumor necrosis factor (TNF) (Genth-Zotz et al. 2002) and have increased thickness of the intestinal wall, elevated intestinal permeability and intestinal ischemia (Sandek et al. 2007). All these observations suggest a causal relationship between HF and gut dysbiosis and the edematous intestinal wall, epithelial dysfunction, and the translocation of LPS and endotoxins through the intestinal epithelial barrier promoting a mechanistic pathway that ultimately aggravates HF and leads to accelerated cardiac decompensation.

Sleep apnea is a highly prevalent comorbidity in HF (Cowie et al. 2017), is characterized by episodic hypoxia and intermittent arousals leading to sleep fragmentation (SF). Like many other disorders, sleep apnea has recently been associated with gut dysbiosis and systemic inflammation (Ko et al, 2019). SF, one of the hallmark components of sleep apnea, has been less extensively examined than intermittent hypoxia (Moreno-Indias et al. 2015; Tripathi et al. 2018), but studies to date have shown that it induces gut dysbiosis (Poroyko et al. 2016), and such changes are reflected by an increase in the Firmicutes/Bacteroidetes ratio, a preferential growth of the families *Lachnospiraceae* and *Rumninococcaceae*, and a decrease in *Lactobacillaceae* (Poroyko et al. 2016). These changes are in turn associated with increased gut permeability, increased systemic LPS levels, and ultimately with systemic inflammation, which can further precipitate and maintain gut dysbiosis (Farre et al, 2018).

Given that both HF and SF are associated with gut dysbiosis and increased inflammation (Farre et al. 2018), we hypothesized that the coexistence of both conditions would result in a more marked alteration of the gut microbiome as compared with either condition in isolation. To test this hypothesis, we analyzed changes in the gut microbiome using a mouse model of HF and SF.

## RESULTS

### Characterization of the microbiome

We used a 16S metabarcoding approach of the V3-V4 region and a computational pipeline (see Materials and Methods) to assess the microbiome composition before and after the treatment, in the different groups. The number of reads observed in each sample ranged from 25,053 to 121,981 with a mean of 58,030.99 (Rarefaction curve, Figure S1. Supplementary material). Overall, we identified 128 and 114 different taxa at the genus and species levels, respectively. We classified 56.76% reads at the genus level, and the five most abundant genera were *Akkermansia, Alistipes, Bacteroides*, *Lachnospiraceae*_NK4136_group and an unclassified *Muribaculaceae* (F.Muribaculaceae.UCG).

We produced Multidimensional scaling (MDS) plots based on the calculated beta diversity (Figure 1). We observed that sample stratification was significantly driven by *Time* (P<0.05 Adonis, in all distance metrics except VAW_GUNifrac). This finding suggests that the microbiota of both treated and control mice had evolved significantly during the four weeks of the experiment (Figure 1A). In addition, we observed that samples clustered in two main enterotypes (Costea et al., 2018) (Figure 1B), which showed a significant relationship with the *Time* variable according to Bray-Curtis dissimilarity (Chi-square, P = 3.228e-06).

**Figure 1.**
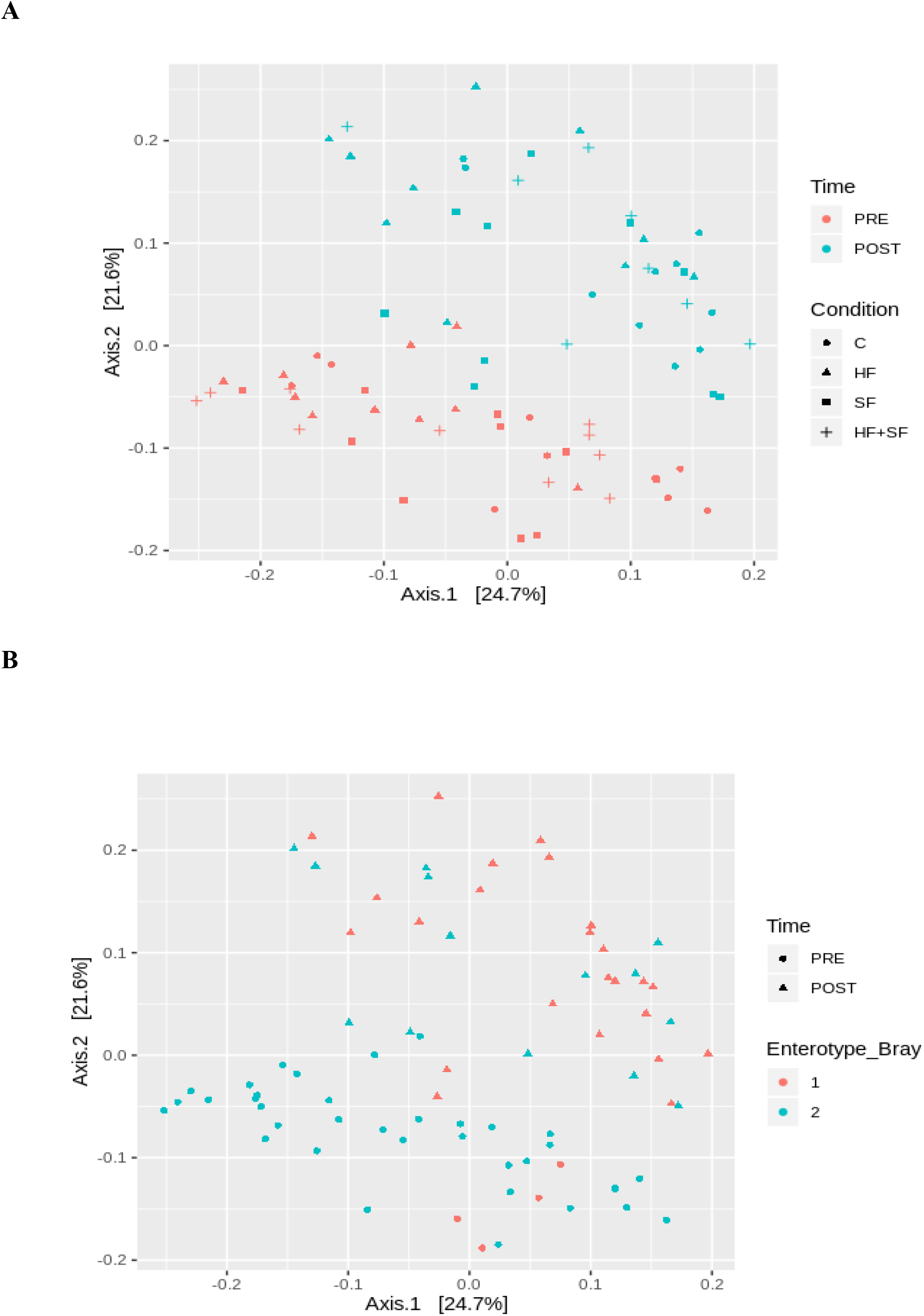
Stratification of the samples. MDS plots based on Bray distance dissimilarity. A) The samples are colored according to the *Time* and shaped according to *Condition* variable B) The samples are colored according to the *Enterotype* variable calculated according to the Bray-Curtis dissimilarity and shaped according to the *Time* variable.

### Alpha diversity

When considering all the samples together, the alpha diversity showed a tendency to increase at the end of the experiment (Figure 2A), although not significantly (P > 0.05, Wilcoxon). However, when comparing alpha diversity before and after the treatment within each group, the control group (C) but not the others, had a significant increase in alpha diversity (Figure 2B), whereas a trend toward a decrease in alpha diversity was noted for HF.

**Figure 2.**
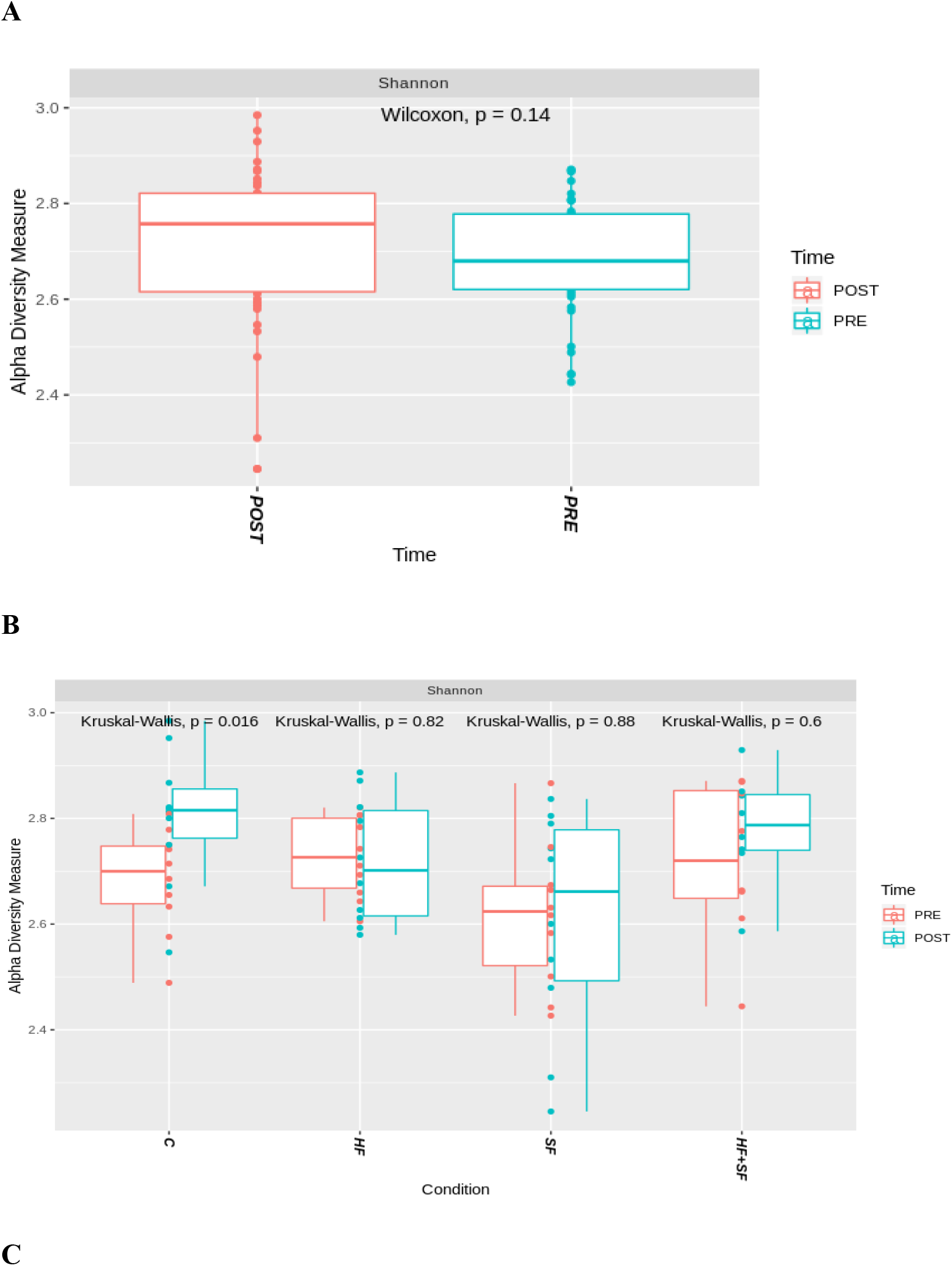
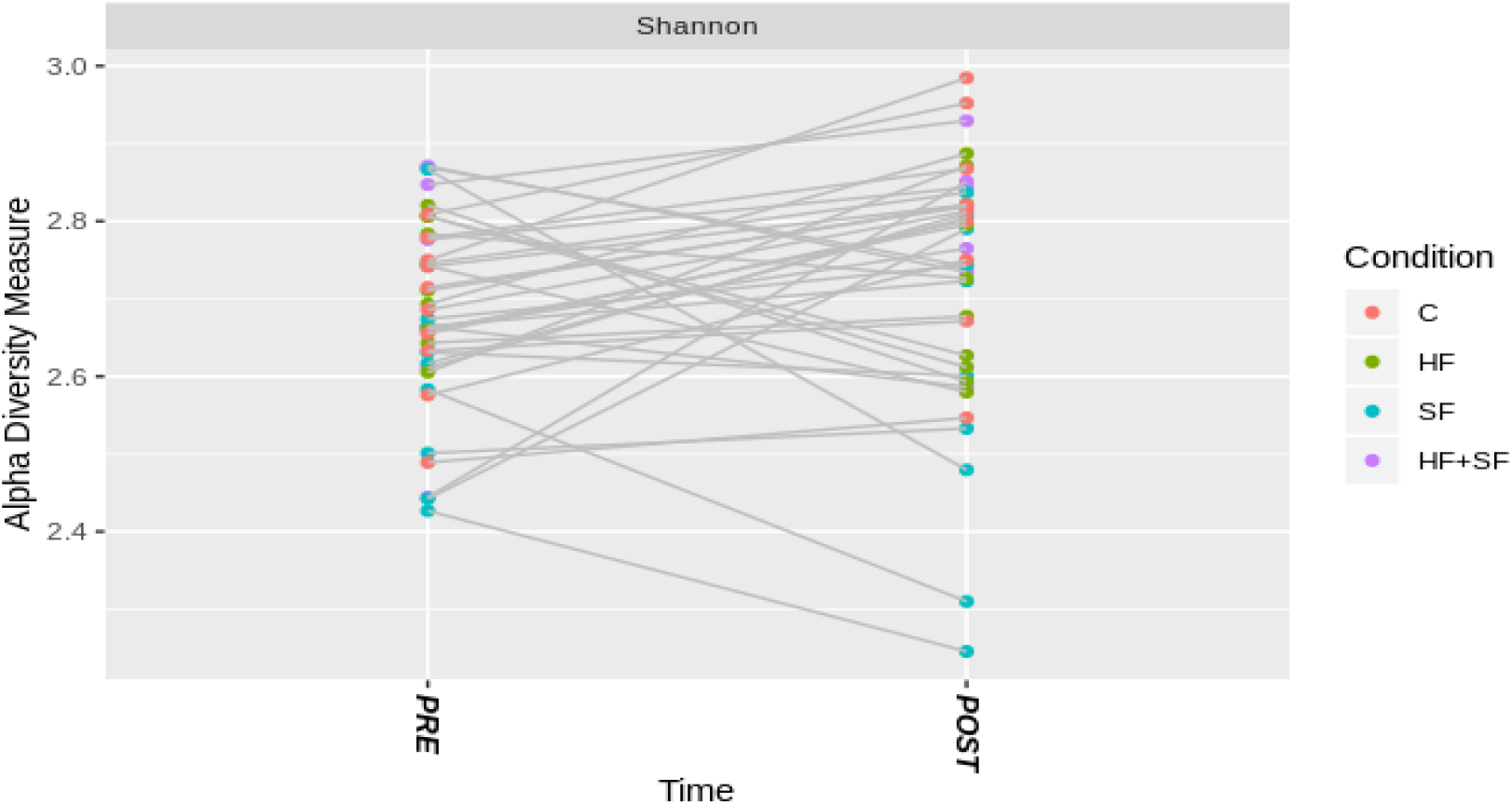
Shannon alpha Diversity measure representation for the paired samples. A) Shannon index according to the *Time* variable B) Shannon index according to the *Condition* variable (C: Controls; HF: Heart Failure; SF: Sleep Fragmentation; HF+SF: Heart Failure and Sleep Fragmentation. C) Variation of Shannon diversity indexes before and after the experiment in each individual mouse. Samples are colored according to the experimental condition.

We also observed differences in alpha diversity between mice subjected to the different conditions. When considering only the samples after the experiment, we observed that both HF and SF groups had significantly lower alpha diversity, as compared to animals in C and (HF+SF) conditions (Figure 3A). When considering all samples, SF mice also showed a significantly lower alpha diversity as compared to the other groups (Figure 3B). This indicates the existence of differences in the basal microbiota before the start of the experiment and highlights the need to focus on changes occurring during the experiment rather than simply comparing final states.

**Figure 3.**
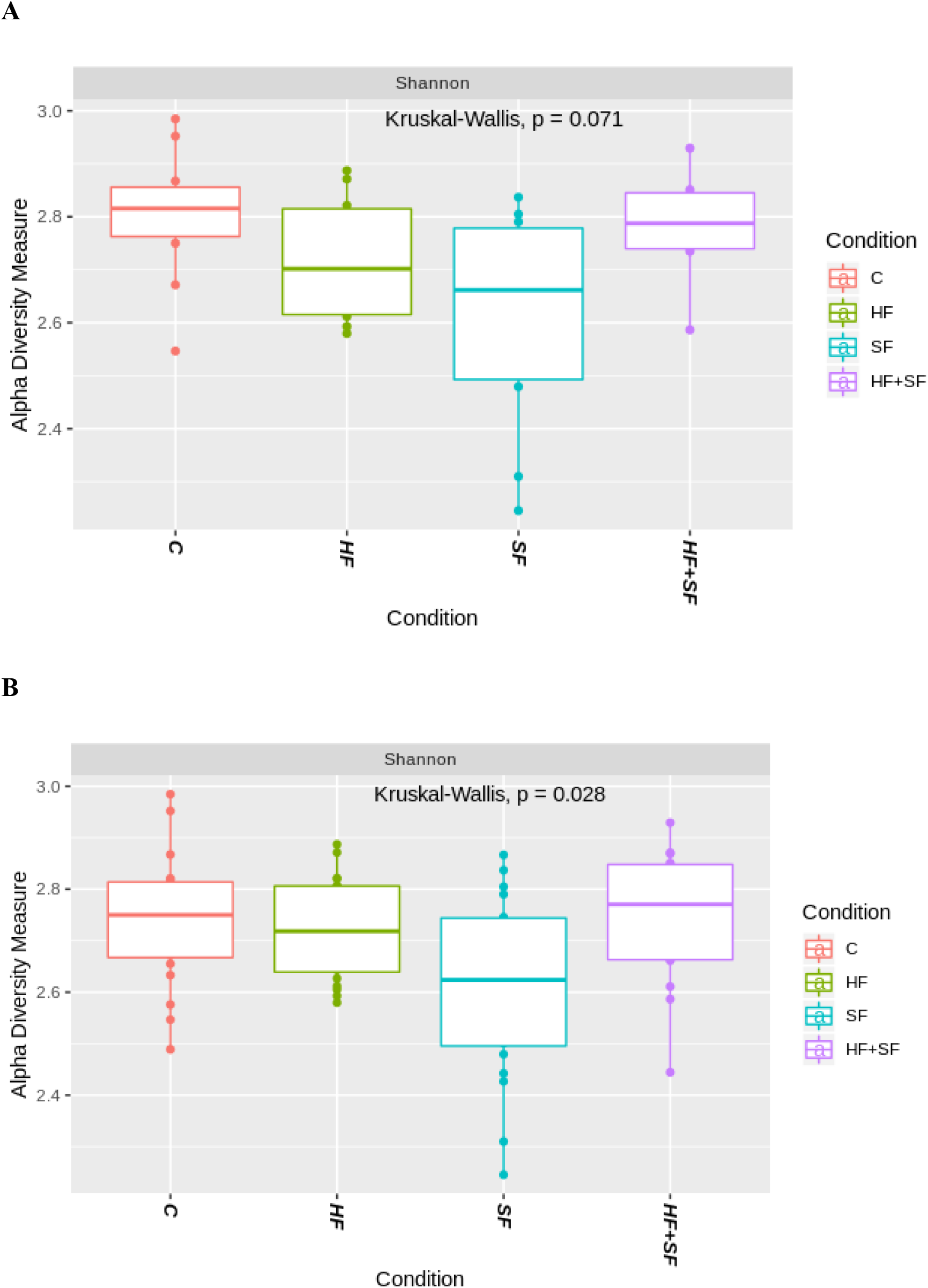
Shannon index representation of the paired samples according to the *Condition v*ariable. The line inside the boxplot represents the median for each of the groups. A) Considering only post samples B) Considering both pre and post samples. Kruskal-Wallis test showed significance (P = 0.028).

### Changes in microbial composition

We observed particular differences in abundance at different taxonomic levels according to the fixed effect variables used in the two different linear models: In the first linear model, all the samples were included and we studied the effect of both the *Condition* and *Time* variables, whereas in the second linear model we included only the samples after the experiment, and focused on the *Condition* and *Change of weight* variables (Table 1).

**Table 1.** Differential abundance analysis findings. A) Linear model including all the samples; Fixed effects: *Condition* and *Time* variable. Random effects: *Batch DNA extraction* and *Animal* (to indicate a paired analysis). B) Linear model taking into consideration only post samples; Fixed effects: *Condition* and *Change of weight* variables. Random effect: *Batch DNA extraction*.

| Variable \ Rank | Phylum | Class | Order | Family | Genus | Species |
| --- | --- | --- | --- | --- | --- | --- |
| Condition | 3 | 5 | 5 | 10 | 23 | 26 |
| Time | 4 | 9 | 10 | 19 | 41 | 47 |

B)
| Variable \ Rank | Phylum | Class | Order | Family | Genus | Species |
| --- | --- | --- | --- | --- | --- | --- |
| Condition | 1 | 2 | 4 | 14 | 30 | 32 |
| Weight change | 1 | 1 | 1 | 3 | 9 | 9 |

For instance, according to the first linear model we obtained 47 differential taxa at the species level according to the *Time* variable. From these taxa, 11 were differentially abundant according to both the *Time* and *Condition* variables: *Bacteroides acidifaciens, Bifidobacterium* spp., F.Atopobiaceae.UCS, *Bacteroides* spp., *Rikenellaceae_RC9_gut_group* spp., F.Lachnospiraceae.UCS, *Ruminococcaceae_UCG.014* spp., *Ruminococcus* spp., *Allobaculum* spp., *Dubosiella* spp. and *Faecalibaculum* spp., whereas 15 and 36 taxa were exclusively reported for *Condition* and *Time* separately, respectively. (Supplementary material, Table 1).

On the other hand, applying the second linear model which only considered post-exposure samples, we observed 32 significantly differentially abundant species according to the *Condition* variable. Applying a multiple comparison test, the comparison with more differences was C versus HF (Figure 4 and Supplementary material, Table 2). Notice that we observed more changes when comparing HF and SF to healthy controls separately instead of when mice were exposed to both conditions. This supports the above mentioned results, in which the alpha diversity was lower in HF or SF separately when compared to either C or HF+SF.

**Figure 4.**
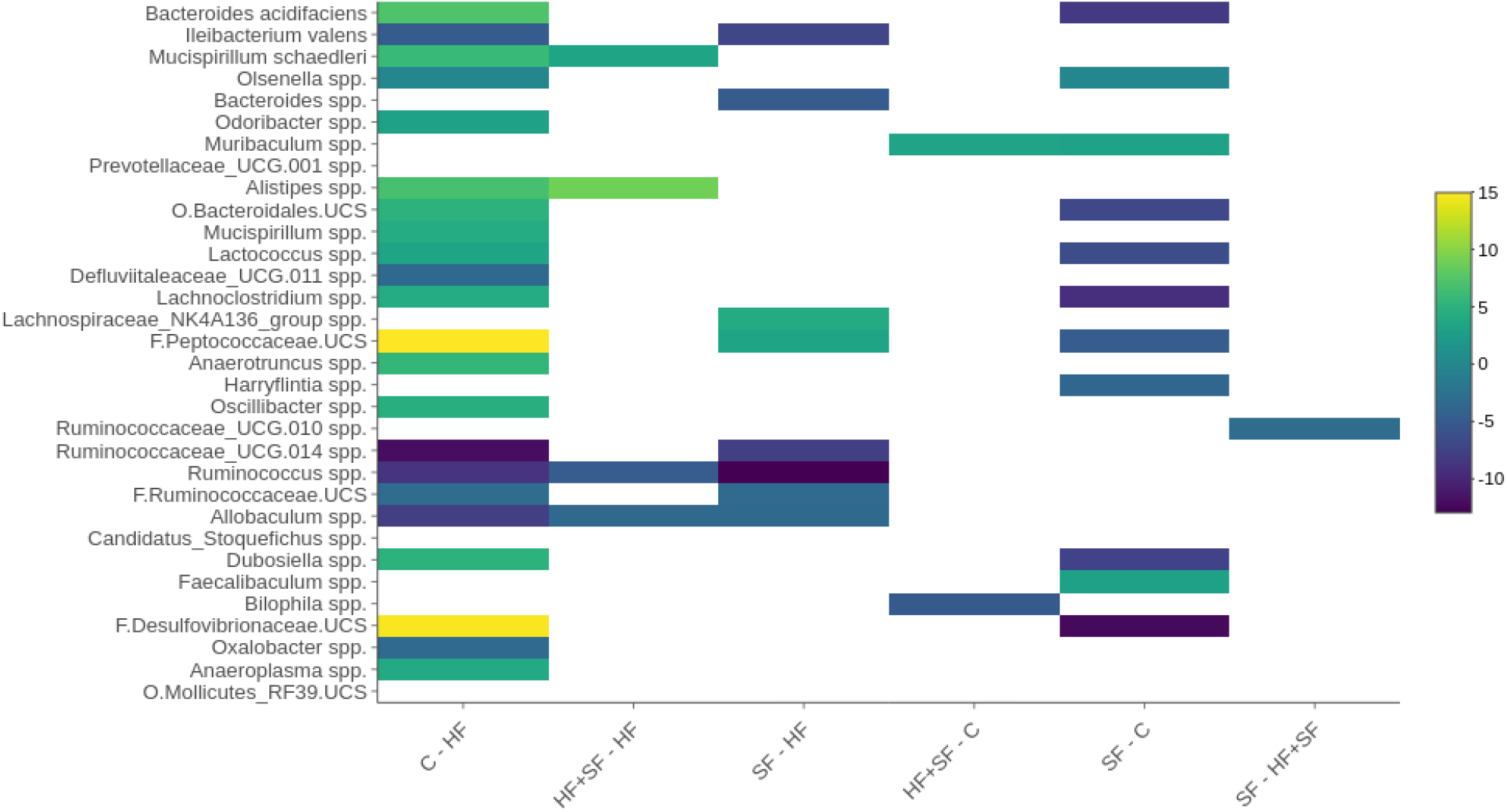
Heatmap representing the 32 significantly differentially abundant taxa at the species level between groups in post samples. The logarithm of only the significant p-values are reported (P < 0.05), where the infinite values are represented as 2.2e-16. The sign of the values was transformed to positive or negative according to the direction of the alteration: positive values for increases in the first group within the comparison and negative values for the decreases. Example: A value of 7.218 for *Bacteroides acidifaciens* when comparing C to HF means that this species is significantly higher in C compared to HF.

Six taxa at the species level were significantly altered by both the *Condition* and *Change of weight* variables: *Ileibacterium valens, Mucispirillum schaedleri*, F.Peptococcaceae.UCS, *Anaerotruncus* spp., *Ruminococcus* spp. and *Allobaculum* spp., while 26 taxa were only significantly differentially abundant according to the *Condition* variable (Table 2).

**Table 2.**
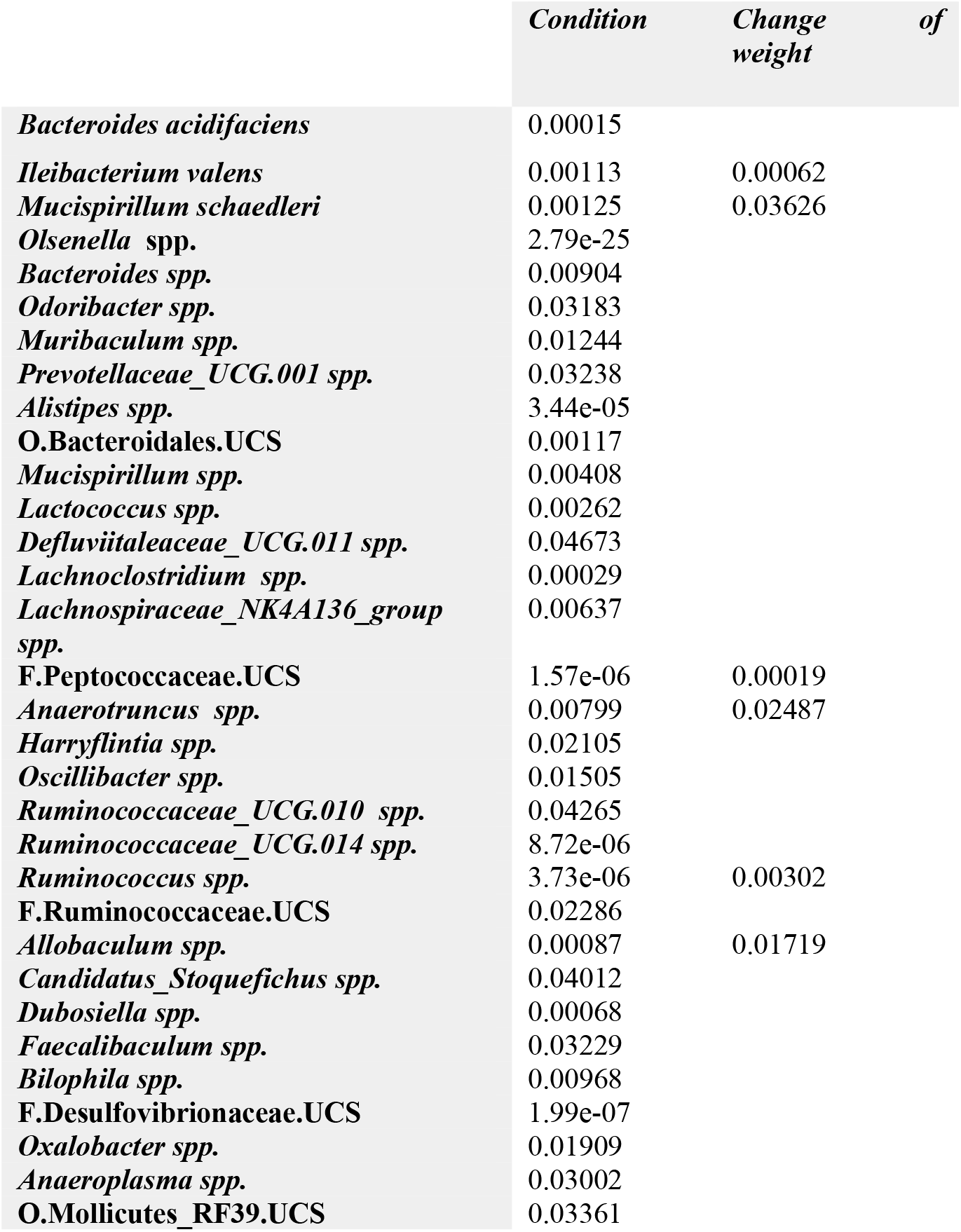
Summary of the p-values corresponding to the 32 significantly differentially abundant taxa at species level according to both *Condition* and *Change of weight* variables.

## DISCUSSION

In the present study we used a mouse model to assess the impact on the gut microbiome composition under conditions of HF and SF, and the combination of the two perturbations, which is frequently present in patients suffering from heart failure who go on to manifest sleep apnea. Overall, the study presents a clear separation between the samples before and after the induction of the conditions, including among the mice in the control group. This clustering may be produced by the anticipated evolution of the microbiome over time, a phenomenon that has been reported in several other studies of the mouse gut microbiome (Kim et al., 2019). Interestingly, an increase in the abundance of the family *Rikenellaceae*, including the genus *Alistipes* (p-value 1.86e-09) in the post group samples (after four weeks of experiment) emerged, taxa that have been previously reported as being overrepresented in old mice and in elderly humans (Langille et al., 2014), (Claesson et al., 2012).

The overall alpha diversity was increased in the post-exposure samples, but this finding was only statistically significant in the control group. This suggests that species richness is significantly higher after the four weeks of the experiment when the mice are allowed to maintain their normal activities and are void of any of the experimental exposures, thereby corroborating earlier studies showing that older individuals exhibit more species overall than juveniles (Mika et al., 2015). These results support the notion of an evolving gut microbiome during mouse development and underscore the importance of including samples taken at the start and at the end of the experiments to control for that variation. Importantly, the variation in species richness differed among the treated groups, wherein those exposed to only one of the relevant conditions displayed diminished species richness. Our findings concur with previous studies that showed an alteration in the microbiome in both HF and SF conditions and a decreased alpha diversity in HF patients (Luedde et al., 2017), (Yuzefpolskaya et al., 2020).

The alteration of both *Lachnospiraceae* and *Ruminococcaceae* observed herein has also been noted by others in both isolated HF or SF models (Luedde et al., 2017; Poroyko et al., 2016). As mentioned, when applying a multiple comparison test considering only post samples, the largest differences were between C and HF. One example of a species that is altered is *Bacteroides acidifaciens*, which decreased in HF compared to C. *B. acidifaciens* has been linked to decreased obesity and to improve insulin sensitivity (Yang et al., 2017), is more abundant in individuals with high-fiber diets and acetate supplementation, and has been reported to play a role in the regulation of the circadian cycle in the heart (Marques et al., 2017; Yang et al., 2017). Since a disturbance in the circadian cycle can cause cardiovascular complications (Duong et al. 2019, Zhang et al. 2020), a decrease in *B. acidifaciens* may serve as an indicator of increased risk for deterioration of the underlying cardiac insufficiency. Interestingly, we also found this species to be decreased in SF samples compared to controls (p-value 0.00025). This could also be due to the same reason, since a disturbed circadian cycle can lead to fragmented sleep, or alternatively, SF could induce the changes in gut microbiome that then disrupt the circadian cycle and elicit increased risk for cardiac decompensation in HF.

When we restrict our attention to the HF models, we observed an increase in the species *Ileibacterium valens* and the genera *Defluviitaleaceae_UCG.011, Ruminococcaceae_UCG.014, Ruminococcus, Allobaculum* and *Oxalobacter* compared to healthy controls. On the other hand, in addition to the mentioned increase of *B. acidifaciens*, we also observed a decrease in the species *Mucispirillum schaedleri* and the genera *Odoribacter, Alistipes, Mucispirillum, Lactococcus, Lachnoclostridium, Anaerotruncus, Oscillibacter, Dubosiella* and *Anaeroplasma*. In previous studies, *Ruminococcaceae_UCG.014* abundance was found as significantly positively associated with serum trimethylamine N-oxide (TMAO) levels, which were associated with coronary atherosclerotic plaque and increased cardiovascular disease risk (Gao et al., 2020). The genus *Ruminococcus* was also found increased in HF models (Cui et al., 2018), and was related to the inflammation that is observed in HF patients by the disruption of the gut barrier through the translocation of gut bacterial DNA and/or endotoxins into the bloodstream (Lataro et al., 2019). It is known that both a high-fat diet (calorie-dense obesogenic) and aging cause inflammation in HF through an alteration of the microbiome such as increasing the phylum Firmicutes, specifically the genus *Allobaculum* (Kain et al., 2019), which in our study was found as significantly more abundant in HF than in C. Both *Alistipes* and *Oscillibacter* were also reported in previous studies as decreased in chronic HF patients (Cui et al., 2018).

Regarding the SF models, we observed increased *Muribaculum* and *Faecalibaculum* at the genus level, and decreased *B. acidifaciens* at the species level and *Lactococcus, Lachnoclostridium, Harryflintia* and *Dubosiella* at the genus level. It is known that melatonin plays a beneficial role in the stabilization of the circadian rhythm (Turek & Gillette, 2004) and a recent study reported that melatonin inhibits *Faecalibaculum* (Hong et al., 2020; Turek & Gillette, 2004). In our study we observed an increase of this genus. Therefore, this reduction can be an indicator of reduced melatonin bioavailability, and consequently reflect a destabilization of the circadian rhythm in SF-exposed mice. Our results also support past findings, whereby the genus *Lachnoclostridium* was reported as underrepresented in chronic intermittent hypoxia in guinea-pigs (Lucking et al., 2018). Hypoxia can be a consequence of a sleep disorder such as sleep apnea. We also found in the bibliography that *Harryflintia* was positively associated with a circadian clock gene (Cry1) whose mutations were related to sleep disorders (Patke et al., 2017).

When considering the coexistence of both HF and SF conditions compared with control mice, we detected only a very small number of differences, namely an increase of *Muribaculum* and a decrease of *Bilophila*. Neither of these genera was previously related to these conditions. Overall, contrary to our initial hypothesis, our results show no strong synergism between the HF and SF conditions as their individual effects were not potentiated when applied in combination. Rather, the changes when the two conditions were combined were less apparent than when applying each condition individually, both in terms of changes in the alpha diversity and in the number of altered taxa. This suggests some level of antagonism between the two conditions, which may influence the microbiome in opposite directions, resulting in some of these effects cancelling each other out.

## CONCLUSION

In summary, we have shown that the gut microbiome contains potential markers of heart failure and of sleep fragmentation when these conditions are evaluated separately. The inflammation observed in HF and SF could be mediated by alterations in abundance of particular taxa. Finally, when the two conditions were applied concomitantly, the alterations in the gut microbiome were milder and virtually disappeared, suggesting some level of antagonism between the effects for HF and SF.

## MATERIALS AND METHODS

### Animal models experiments

Forty male mice (C57BL/6J; 10 weeks old; 12 h light/dark cycle; water/food *ad libitum*) were randomly allocated into four groups (n=10 each). In two groups, the mice were allowed to sleep normally: healthy control (C) and heart failure (HF). In two groups (SF, HF+SF), SF was imposed, and in two groups (HF, HF+SF) heart failure was induced. The animal experiment including the setting of the HF and SF models were approved by the institution ethical committee and has been recently described in detail (Cabrera-Aguilera et al, 2020).

HF was induced by continuous infusion of isoproterenol (Cabrera-Aguilera et al, 2020). Briefly, mice were anesthetized by isoflurane inhalation and an osmotic minipump (Alzet, model 1004) was implanted subcutaneously in the flank. The pump delivered 30 mg/kg per day of isoproterenol (Sigma Aldrich; in sterile 0.9% NaCl solution) for 28 days. Buprenorphine (0.3 mg/kg, i.p.) was administered 10 minutes before surgery and after 24 hours, and the suture was removed 7 days after surgery. Healthy animals were subjected to the same protocol with the only difference being that no isoproterenol was dissolved into the 0.9% NaCl pump medium. As described elsewhere (Cabrera-Aguilera et al, 2020), the effectiveness of the HF model in these animals was assessed by echocardiography after 28 days of isoproterenol infusion, confirming that mice in the HF groups had significant increases in left ventricular end-diastolic and LVESD and end-systolic diameter as well as significant reductions in left ventricular ejection fraction and fraction shortening.

Two days after surgery, SF was induced daily by means of a previously described and validated device for mice (Lafayette Instruments, Lafayette, IN), which is based on intermittent tactile stimulation with no human intervention. Sleep arousals were induced by a mechanical near-silent motor with a horizontal bar sweeping just above the cage floor from one side to the other side in the standard mouse laboratory cage. Each sweep was applied in 2-minute intervals during the murine sleep period (8 a.m. to 8 p.m.) for 28 days (until day 30 from surgery) (Cabrera-Aguilera et al, 2020).

At the end of the 4-week experiment (HF, SF, HF+SF and control), fecal samples were obtained directly from stool expulsion stimulated by manual handling of the animal and were immediately frozen at −80°C and stored until analyzed.

### DNA extraction, library preparation and sequencing

DNA was extracted from mice fecal individual samples using the DNeasy PowerLyzer PowerSoil Kit (Qiagen, ref. QIA12855) following the manufacturer’s instructions. After adding mice stool samples to the PowerBead Tubes, 750 μl of PowerBead Solution and 60 μl of Solution C1 were added, and samples were vortexed briefly and incubated at 70°C with shaking (700 rpm) for 10 min. The extraction tubes were then agitated twice in a 96-well plate using Tissue lyser II (Qiagen) at 30 Hz/s for 5 min.

Tubes were centrifuged at 10,000 g for 3 min and the supernatant was transferred to a clean tube. 250 μl of Solution C2 were added, and samples were vortexed for 5 s and incubated on ice for 10 min. After 1 min centrifugation at 10,000 g, the supernatant was transferred to a clean tube, 200 μl of Solution C3 were added, and samples were vortexed for 5 s and incubated on ice for 10 min again. 750 μl of the supernatant were transferred into a clean tube after 1 min centrifugation at 10,000 g. Then, 1,200 μl of Solution C4 were added to the supernatant, samples were mixed by pipetting up and down, and 675 μl were loaded onto a spin column and centrifuge at 10,000 g for 1 min, discarding the flow through. This step was repeated three times until all samples had passed through the column. 500 μl of Solution C5 were added onto the column and samples were centrifuged at 10,000 g for 1 min, the flow through was discarded and one extra minute centrifugation at 10,000 g was done to dry the column. Finally, the column was placed into a new 2 ml tube to the final elution with 50 μl of Solution C6 and centrifugation at 10,000 g for 30 s.

Four μl of each DNA sample were used to amplify the V3–V4 regions of the bacterial 16S ribosomal RNA gene, using the following universal primers in a limited cycle PCR:

V3-V4-Forward (5-TCGTCGGCAGCGTCAGATGTGTATAAGAGACAGCCTACGGGNGGCWGCAG-3’) and V3-V4-Reverse (5-GTCTCGTGGGCTCGGAGATGTGTATAAGAGACAGGACTACHVGGGTATCTAAT CC-3’).

To prevent unbalanced base composition in further MiSeq sequencing, we shifted sequencing phases by adding various numbers of bases (from 0 to 3) as spacers to both forward and reverse primers (we used a total of 4 forward and 4 reverse primers). The PCR was performed in 10 μl volume reactions with 0.2 μM primer concentration and using the Kapa HiFi HotStart Ready Mix (Roche, ref. KK2602). Cycling conditions were initial denaturation of 3 min at 95 °C followed by 20 cycles of 95 °C for 30 s, 55 °C for 30 s, and 72 °C for 30 s, ending with a final elongation step of 5 min at 72 °C.

After the first PCR step, water was added to a total volume of 50 μl and reactions were purified using AMPure XP beads (Beckman Coulter) with a 0.9X ratio according to manufacturer’s instructions. PCR products were eluted from the magnetic beads with 32 μl of Buffer EB (Qiagen) and 30 μl of the eluate were transferred to a fresh 96-well plate. The primers used in the first PCR contain overhangs allowing the addition of full-length Nextera adapters with barcodes for multiplex sequencing in a second PCR step, resulting in sequencing ready libraries. To this end, 5 μl of the first amplification were used as template for the second PCR with Nextera XT v2 adaptor primers in a final volume of 50 μl using the same PCR mix and thermal profile as for the first PCR but only 8 cycles. After the second PCR, 25 μl of the final product was used for purification and normalization with SequalPrep normalization kit (Invitrogen), according to the manufacturer’s protocol. Libraries were eluted in 20 μl and pooled for sequencing.

Final pools were quantified by qPCR using Kapa library quantification kit for Illumina Platforms (Kapa Biosystems) on an ABI 7900HT real-time cycler (Applied Biosystems). Sequencing was performed in Illumina MiSeq with 2 × 300 bp reads using v3 chemistry with a loading concentration of 18 pM. To increase the diversity of the sequences 10% of PhIX control libraries were spiked in.

Two bacterial mock communities were obtained from the BEI Resources of the Human Microbiome Project (HM-276D and HM-277D), each contained genomic DNA of ribosomal operons from 20 bacterial species. Mock DNAs were amplified and sequenced in the same manner as all other murine stool samples. Negative controls of the DNA extraction and PCR amplification steps were also included in parallel, using the same conditions and reagents. These negative controls provided no visible band or quantifiable DNA amounts by Bioanalyzer, whereas all of our samples provided clearly visible bands after 20 cycles.

### Microbiome analysis

The *dada2* pipeline (v. 1.10.1) (Callahan et al., 2016) was used to obtain an ASV (amplicon sequence variants) table (Nearing et al., 2018). First, the sequence quality profiles of forward and reverse sequencing reads were examined using the *plotQualityProfile* function of dada2. Based on these profiles, low-quality sequencing reads were filtered out and the remaining reads were trimmed at positions 285 (forward) and 240 (reverse). The first 10 nucleotides corresponding to the adaptors were also trimmed, using the *filterAndTrim* function with the following parameters:

> “filterAndTrim(fnFs, filtFs, fnRs, filtRs, truncLen=c(285,240), maxN=0,
>
> maxEE=c(10,10), truncQ=1, rm.phix=TRUE, trimLeft=c(10,10), compress=TRUE, multithread=TRUE)”

Then, identical sequencing reads were combined into unique sequences to avoid redundant comparisons (dereplication), sample sequences were inferred (from a pre-calculated matrix of estimated learning error rates) and paired reads were merged to obtain full denoised sequences. From these, chimeric sequences were removed. Taxonomy was assigned to ASVs using the *SILVA* 16s rRNA database (v. 132) (Quast et al., 2013). Next, a phylogenetic tree representing the taxa found in the sample dataset was reconstructed by using the phangorn (v. 2.5.5) (Schliep, 2011) and Decipher R packages (v 2.10.2) (Wright et al., 2016). We integrated the information from the ASV table, Taxonomy table, phylogenetic tree and metadata (information relative to the samples such as the time, batch of the DNA extraction and change of weight) to create a *phyloseq* (v. 1.26.1) object (McMurdie & Holmes, 2013). Positive and negative sequencing controls (mock communities and water samples, respectively) sequenced and included in the ASV table were removed from subsequent statistical analyses.

The metadata consisted of 11 variables: *batchDNAextraction, sample, Time* (indicating whether samples were taken prior to or post treatment); *Box; SF.NORMAL.SLEEP* (Sleep fragmentation or normal sleep); *Animal*; *Pump* (What substance was injected, Isoproterenol or Saline - control); *Initial_weight; Final_weight*; and *Initial ecography* (the value of which was “Ready” for all the animals). We created a new variable called *Condition* corresponding to the four different treatment groups: C, HF, SF and HF+SF.

Taxonomic composition metrics such as alpha-diversity (within-sample) and beta-diversity (between samples) were characterized. Using the *estimate_richness* function of the *phyloseq* package we calculated the alpha diversity metrics including Observed.index, Chao1, Shannon, Simpson and InvSimpson indices. Regarding the different beta-diversity metrics, we used the *Phyloseq* and *Vegan* (v. 2.5-6) (Oksanen et al. 2019) packages to characterize nine distances based on differences in taxonomic composition of the samples including JSD, Weighted-Unifrac, Unweighted-unifrac, VAW-Gunifrac, a0-Gunifrac, a05_Gunifrac, Bray, Jaccard and Canberra. We also computed Aitchison distance (Gloor et al., 2017) using the *cmultRepl* and *codaSeq.clr* functions from the *CodaSeq* (v. 0.99.6) (Gloor & Reid, 2016) and *zCompositions* (v.1.3.4) (Palarea-Albaladejo & Martín-Fernández, 2015) packages.

Normalization was performed by transforming the data to relative abundances, and samples containing fewer than 950 reads were discarded and taxa that appeared in fewer than 5% of the samples at low abundances were filtered out:

> “prune_samples(sample_sums(object) >= 950, object)”
>
> “filter_taxa(object, function(x) sum(x > 0.001) > (0.05 * length(x)), prune = TRUE)”

### Statistical analysis

Comparison of echocardiographic data between all groups at baseline was performed using one-way ANOVA. Comparison of echocardiographic data between all groups at day 30 was performed using two-way ANOVA followed by the Student-Newman-Keuls comparison method. The data is presented as mean ± SEM.

We used the Partitioning Around Medoid (PAM) algorithm (Reynolds et al., 2006), as implemented in the *cluster* library (v. 2.0.7-1), to explore clustering of the samples. We further evaluated this, performing a Permutational Multivariate Analysis of Variance (PERMANOVA) using the ten-distance metrics mentioned above, and the *adonis* function from the *Vegan* R package (v. 2.5-6) (Oksanen et al. 2019). The *Time* and *Box* variables were considered as covariates.

To identify taxonomic features (Phylum, Class, Order, Family, Genus and Species) that show significantly different abundances among studied conditions, we used linear models, as implemented in the R package *lme4* (v. 1.1-21) (Bates et al. 2015). Two different linear models were built: In the first one, the fixed effects were the *Condition* and *Time* variables and the random effects were the *batch of the DNA extraction* and the *animal*, where this last one is an indicator of a paired analysis (tax_element ~ Condition + Time + (1| batchDNAextraction) + (1|Animal)). On the other hand, in the second linear model we included only post samples and instead of the *Time* variable, we used as a fixed effect the *Change of weight* of the mouse models (*Final_weight - Initial_weight*). In this case we only used as a random effect the batch (tax_element ~ Condition_POST_only + Change_of_weight + (1|batchDNAextraction)).

Analysis of Variance (ANOVA) was applied to assess the significance for each of the fixed effects included in the models using the *Car* R package (v. 3.0-6) (Fox et al., 2013). To assess particular differences between groups we performed multiple comparisons to the results obtained in the linear models using the *multcomp* R package (v. 1.4-12) (Hothorn et al., 2008). We applied Bonferroni as a multiple testing correction. Statistical significance was defined when p values were lower than 0.05 in all the analyses.

## ACKNOWLEDGMENTS

The authors wish to thank Mrs. Elisabeth Urrea and Mr. Miguel A. Rodriguez-Lazaro for their excellent technical assistance.

## Data availability

Raw sequence data can be found in the Sequence Read Archive with the Bioproject accession code: PRJNA662468

## Authors contribution

O. Khannous-Lleiffe, J.R. Willis and E. Saus carried out the microbiota analysis. I. Cabrera-Aguilera was in charge of the animal model experiments. I. Almendros, R. Farré and D. Gozal participated in data interpretation and scientific discussion. T. Gabaldón designed and supervised the microbiota analysis and discussion. Nuria Farré conceived the study and supervised the whole research. All authors participated in the manuscript preparation.

